# Transcriptome profile and clinical characterization of ICOS expression in brain gliomas

**DOI:** 10.1101/2022.05.17.492394

**Authors:** Jin Wang, Fei Shi, Aijun Shan

## Abstract

**Aims:** Inducible Co-Stimulator (ICOS), an immune costimulatory molecule, has been found to play an essential role across various malignancies. This study investigated the transcriptome profile and clinical characterization of ICOS in gliomas.

**Methods:** Clinical information and transcriptome data of 301 glioma samples were downloaded from the Chinese Glioma Genome Atlas (CGGA) data set for analysis. Furthermore, the results were validated in 697 samples with RNAseq data from the TCGA glioma data set. In addition, single-cell sequencing data from CGGA and GSE 163108 datasets were used to analyze the ICOS expression across different cell types. Statistical analyses and figure production were performed with R-language.

**Results:** We found that ICOS was significantly upregulated in higher-grade, IDH wildtype, and mesenchymal subtype of gliomas. Functional enrichment analyses revealed that ICOS was mainly involved in glioma-related immune response. Moreover, ICOS showed a robust correlation with other immune checkpoints, including PD1/PD-L1/PD-L2 pathway, CTLA4, ICOSL (ICOS ligand), and IDO1. Furthermore, based on seven clusters of metagenes, GSVA identified that ICOS was tightly associated with HCK, LCK, MHC-I, MHC-II, STAT1, and interferon, especially with LCK, suggesting a strong correlation between ICOS and T-cell activity in gliomas. In cell lineage analysis, ICOS-higher gliomas tended to recruit dendritic cells, monocytes, and macrophages into the tumor microenvironment. Single-cell sequencing analysis indicated that ICOS was highly expressed by regulatory T cells (Treg). Finally, patients with higher ICOS had shortened survival. ICOS was an independent prognosticator for glioma patients.

**Conclusions:** Higher ICOS was correlated with more malignancy of gliomas and significantly associated with Treg activity among glioma-related immune responses. Moreover, ICOS could contribute as an independent prognostic factor for gliomas. Our study highlighted the role of ICOS in glioma and may facilitate therapeutic strategies targeting ICOS for glioma.

## 1. Introduction

Brain gliomas, accounting for 70% of primary intracranial tumors in adult patients, are commonly characterized by a high mortality and disability rate(1). Despite substantial improvements in management, the unfavorable outcomes for gliomas remain unchanged, especially for the most aggressive type - GBM(2, 3); the median survival time remains less than 15 months. In the last decades, substantial immunotherapy advancements in other cancers have brought new hope for glioma treatment. Multiple studies have identified immune targets for glioma immunotherapy, mainly focusing on co-inhibitory checkpoints, including PD1/PD-L1 and CTLA4. However, most gliomas failed to respond to current immunotherapy, which facilitated us finding additional immune checkpoints for gliomas.

Inducible Co-stimulator (ICOS, also termed as CD278, H4, AILIM), a member of the costimulatory molecule family consisting of ICOS, CD28, and CTLA4, is abundantly expressed on the cell surface of the activated and mature T-cells rather than naïve T-cells(4, 5). The costimulatory signal is induced after ICOS engagement with its unique ligand (ICOSL) to facilitate a series of immune-related biological processes, including development of germinal centers (6), activation of T-cell-dependent B cells, and switching of antibody class (7). More importantly, the ICOS/ICOSL pathway is particularly essential for T-cells themselves, promoting their differentiation, proliferation, activation, and survival(5, 8). In addition, ICOS enhances the secretion of multiple immune cytokines, including TNF-α, IL-4, IL-5, IL-6, IL-10, and IL-21 (4, 5, 9, 10). The abnormality of ICOS expression leads to a range of pathophysiological dysfunctions, such as immunodeficiency, opportunistic infection, and malignant tumors.

ICOS has been widely reported as an important immune checkpoint among various cancers, including melanoma, gastrointestinal and liver cancer, gynecological cancer, breast cancer, renal clear cell carcinoma, and Merkel carcinoma. However, the roles of ICOS across different types of malignancies are inconsistent, mainly due to the dualistic effect of ICOS in the tumor microenvironment. On the one hand, ICOS exerts its anti-tumor effect through the enhancement of CD4+ and CD8+ effector T cells(11, 12); on the other hand, ICOS significantly activates and upregulates regulatory T cells (Tregs)(13), which are a subpopulation of T-cells that mainly functions as an immunosuppressor and consequently facilitate the immune escape in tumors(14). Therefore, many studies sought to determine the correlation between ICOS and outcomes of malignant tumors, as expected, yielded contradictory results. For patients with melanoma(15-17), gastric(18, 19) and liver cancer(20), gynecological(21, 22) and breast cancer(23, 24), as well as renal clear cell carcinoma(25), higher ICOS expression predicted much worse survival. In contrast, among patients who suffered from colorectal cancer(26, 27) and follicular lymphoma(28), the upregulation of ICOS expression yielded a better prognosis.

As an important immune costimulatory molecule, ICOS has attracted more and more attention in both hematologic and solid malignancies. However, no comprehensive report on gliomas has been reported. Only one study performed by Gousias et al.(29) investigated the fraction of ICOS+ Treg via immune-cell count analysis on 29 glioma patients. To elucidate the ICOS expression profile and clinical characterization in pan-glioma, we collected microarray data of 301 glioma samples from the Chinese Glioma Genome Atlas (CGGA) dataset and performed this integrative analysis of ICOS among whole grade gliomas. Subsequently, the results were further validated in 697 gliomas with RNAseq of TCGA dataset. Our study would be the first comprehensive report demonstrating molecular and clinical characterization of ICOS expression among pan-gliomas. We believe ICOS will become a hotspot for immunotherapy of gliomas.

## 2. Materials and Methods

### 2.1. Sample and data collection

Microarray data and corresponding clinical information of 301 glioma patients (WHO grade II to IV) were obtained from the Chinese Glioma Genome Atlas (CGGA)(30) dataset (http://www.cgga.org.cn/). The validation cohort utilized the TCGA glioma dataset (http://cancergenome.nih.gov/)(31), which included 697 patients (WHO grade II to IV) with RNAseq data (RSEM-normalized, level 3). Totally, 998 glioma patients were included in this study. **Table S1** summarized the baseline characteristics of both cohorts. In addition, the single-cell RNA-seq data (sc-RNAseq) of glioma patients were obtained from CGGA(32), which consisted of 6148 cells collected from 73 regions of 14 patients. Further, the sc-RNAseq dataset of GSE163108(33), consisting of 25256 glioma-infiltrating T cells from 31 adult patients, was downloaded from the GEO website. Of them, there were 3277 CD4+ T cells, 21502 CD8+ T cells, 89 cycling T cells, and 388 regulatory T cells (Treg). The study was approved by the Ethics Committee of Shenzhen People’s Hospital, and written consents were waived due to the use of de-identified patient data from public datasets.

### 2.2. Statistical analysis

For RSEM-normalized RNA sequencing data from TCGA dataset, log2 transformation was performed before analysis, while microarray data from CGGA dataset, which had already been preprocessed by CGGA project, were analyzed directly. Gaussian distribution test was performed before data analysis. For sc-RNAseq data from CGGA and GSE163108, the data provider had already excluded the low-quality genes and low-quality cells. The percentage of mitochondria-expressed genes was less than 5% in CGGA, and 10%in GSE163108, respectively.

ICOS co-expressed genes were identified according to the correlation coefficients of Pearson correlation (CGGA) or Spearman correlation (TCGA). The correlation coefficient |r| > 0.5 was set as the filter criteria to screen out significantly-correlated genes of ICOS in each dataset. Gene Ontology (GO) analyses were performed on DAVID(34) website (2021 update version, https://david.ncifcrf.gov/). Hallmark genesets (h.all.v7.5.1.symbols.gmt) were downloaded from the GSEA(35) website (http://software.broadinstitute.org/). The gene order for GSEA analysis was pre-ranked according to the correlation coefficient value with ICOS. The number of permutations was 1000. Normalized enrichment score (NES) > 1 and *False discovery rate (FDR)* < 0.25 were considered as significantly enriched in the geneset. Seven gene clusters consisting of 104 genes, which represented distinct inflammatory activities, were termed metagenes(36) (**Table S2**), and subsequent Gene Sets Variation Analysis (GSVA)(37) was used for evaluation of the metagene expression level. The immune score, stroma score, microenvironment score, and ICOS-correlated immune cell subpopulations were calculated with the XCELL(38).

R language, together with a series of R-packages, including ggplot2, pheatmap, pROC, circlize, corrgram, clusterprofiler(39), survival, survminer, VennDiagram, as well as forestmodel were utilized for statistical analyses and graphical work. Sc-RNAseq analysis was performed with the Seurat package, and PCA was used for dimensional reduction with the resolution of 0.1 for CGGA and 0.2 for GSE163108. Cox regression analysis was performed with coxph function provided in the R-package of survival. All statistical tests were two-sided. A *p*-value less than 0.05 was considered statistically significant.

## 3. Results

### 3.1. ICOS is associated with a more malignant phenotype of glioma

ICOS expression levels among the WHO grades were compared. Similar results were obtained from both CGGA and TCGA datasets and revealed that a higher grade was usually paralleled with an increased ICOS expression. Though no significant difference was detected between grades III and IV in CGGA dataset, an increasing trend of ICOS could also be observed (**Figures. 1A and 1E**). These findings suggested that ICOS upregulation was associated with more malignancy in gliomas. Moreover, IDH-wildtype gliomas were found to be associated with a higher ICOS expression pattern compared to IDH-mutant counterparts in both datasets (Figures. 1B and 1F), which further confirmed the correlation between ICOS and the aggressiveness in gliomas. Furthermore, ICOS expression was compared across different molecular subtypes. As shown in **Figures 1C and 1G**, ICOS was significantly upregulated in the mesenchymal than that in other subtypes, suggesting the potential discriminatory power of ICOS for mesenchymal-subtype gliomas. ROC curves showed that AUCs were 73.7% (**Figure. 1D**) in CGGA and 85.1% (**Figure. 1H**) in TCGA. These results suggested that ICOS was inclined to express more in higher-grade gliomas, IDH-wildtype, and mesenchymal subtype. ICOS might exert a pro-tumoral effect on tumorigenesis and development of gliomas.

**Figure 1.**
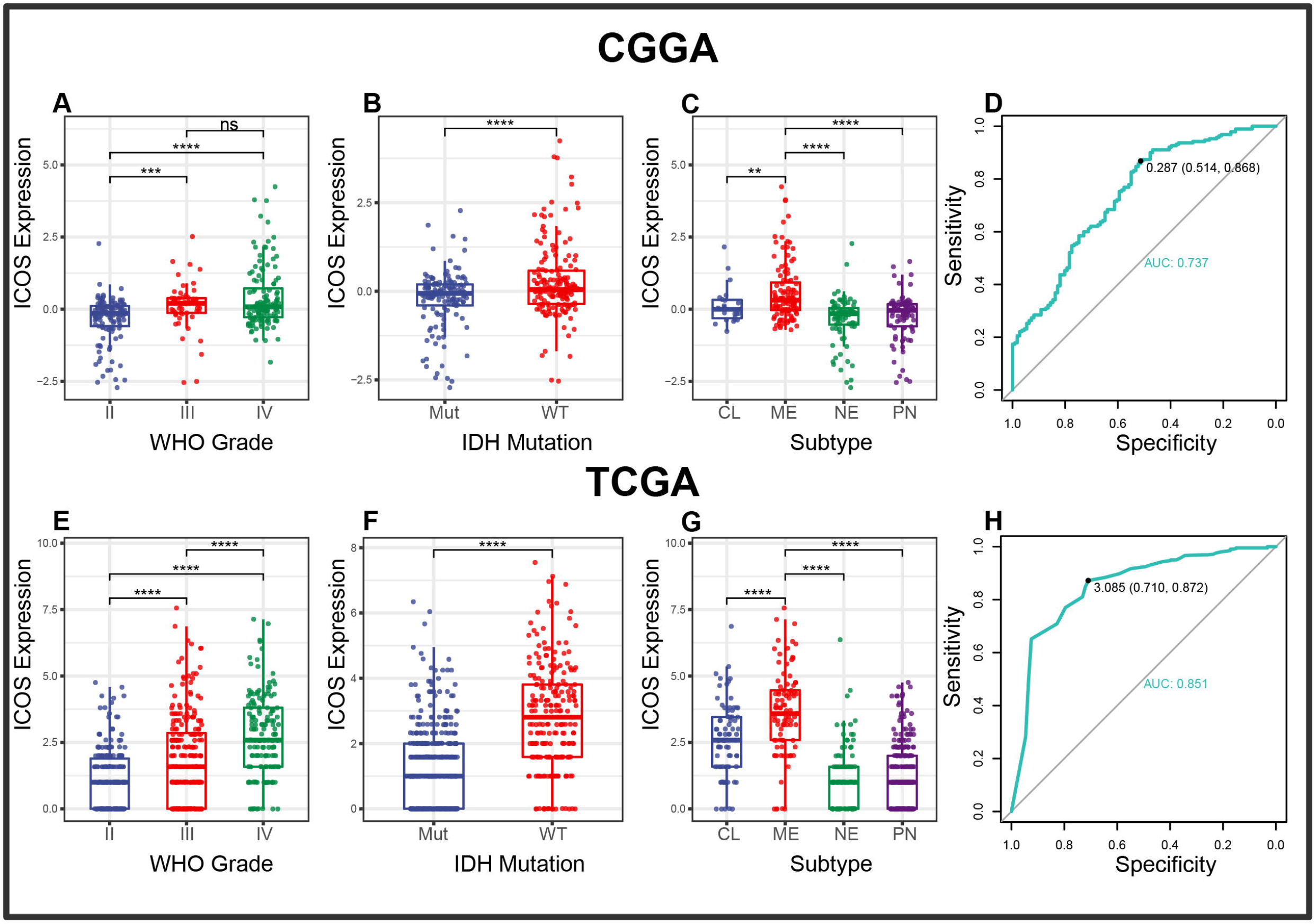
ICOS expression in CGGA and TCGA dataset according to WHO grade (A), IDH mutation status (B), TCGA molecular subtype (C) and ROC curves (D) for distinguishing mesenchymal subtype. * indicates p value < 0.05, **indicates p value < 0.01, *** indicates p value < 0.001, **** indicates p value < 0.0001.

### 3.2. ICOS is involved in glioma-related immune response

To identify the biological functions related to ICOS, ICOS co-expressed genes were screened out for GO analysis. In the CGGA dataset, 1050 ICOS co-expressed genes, consisting of 945 positively correlated and 105 negatively correlated, were identified. It turned out that genes positively correlated with ICOS were mainly involved in immune response, while genes that were negatively correlated with ICOS were associated with a normal biological function (synapse organization) (**Figure 2A**). In the TCGA dataset, 783 ICOS co-expressed genes, all positively correlated with ICOS, were also involved in a range of immune-related biological processes. To ensure an accurate result, ICOS co-expressed genes overlapped between CGGA and TCGA were further filtered for functional annotation, and we found that ICOS-related genes were mainly associated with T cell activation. **(Figures. S2A and S2B)**. Given the distinctive property of GBM, we further evaluated ICOS-related biological processes in GBM. Significantly-correlated genes of both cohorts were further annotated, and we found that ICOS showed an even higher correlation with immune response and T cell activation (**Figures 2C, 2D, S2C, and S2D**). These results indicated that ICOS was mainly involved in glioma-related immune response.

**Figure 2.**
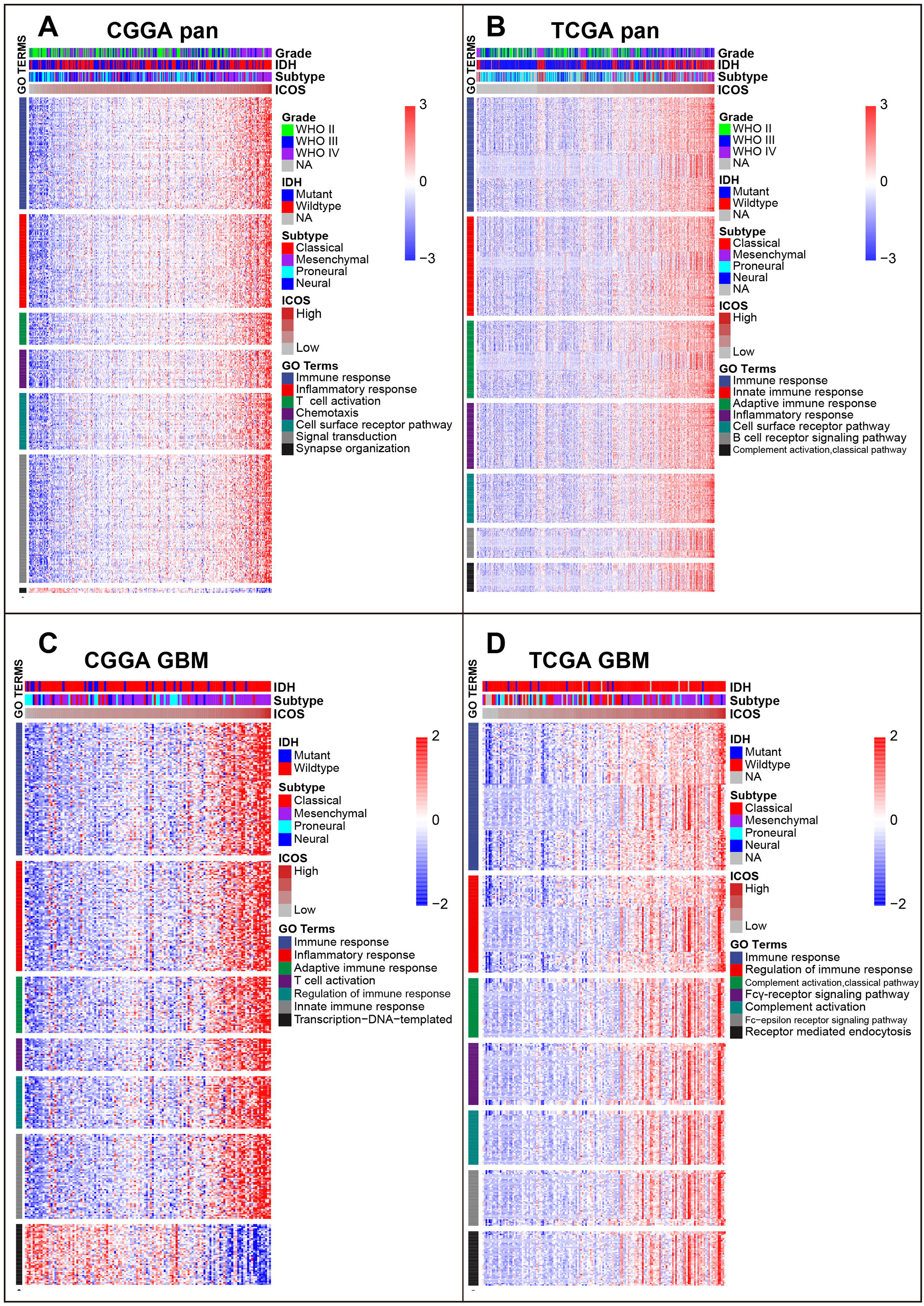
Gene ontology analysis for ICOS in pan-glioma (A, B) and glioblastoma (C, D)

GSEA analyses using the Hallmark genesets were further performed to validate the ICOS-related biological process. ICOS showed robust positive correlation with HALLMARK ALLOGRAFT REJECTION in CGGA (NES = 2.872, *FDR* < 0.001) (**Figure 3A**). followed by other Hallmark genesets, including interferon-gamma response, inflammatory response, interferon-alpha response, and IL6-JAK-STAT3 signaling, which all strongly pointed to the participation of ICOS in the glioma-related immune response. The results were further validated in TCGA (HALLMARK ALLOGRAFT REJECTION, NES = 3.528, *FDR* < 0.001) **(Figure 3B)**. Moreover, we observed a similar pattern of GSEA results in GBM of both cohorts **(Figures 3C and 3D)**, further confirming the profound association of ICOS with glioma-related immune response, consistent with what we observed in GO analysis.

**Figure 3.**
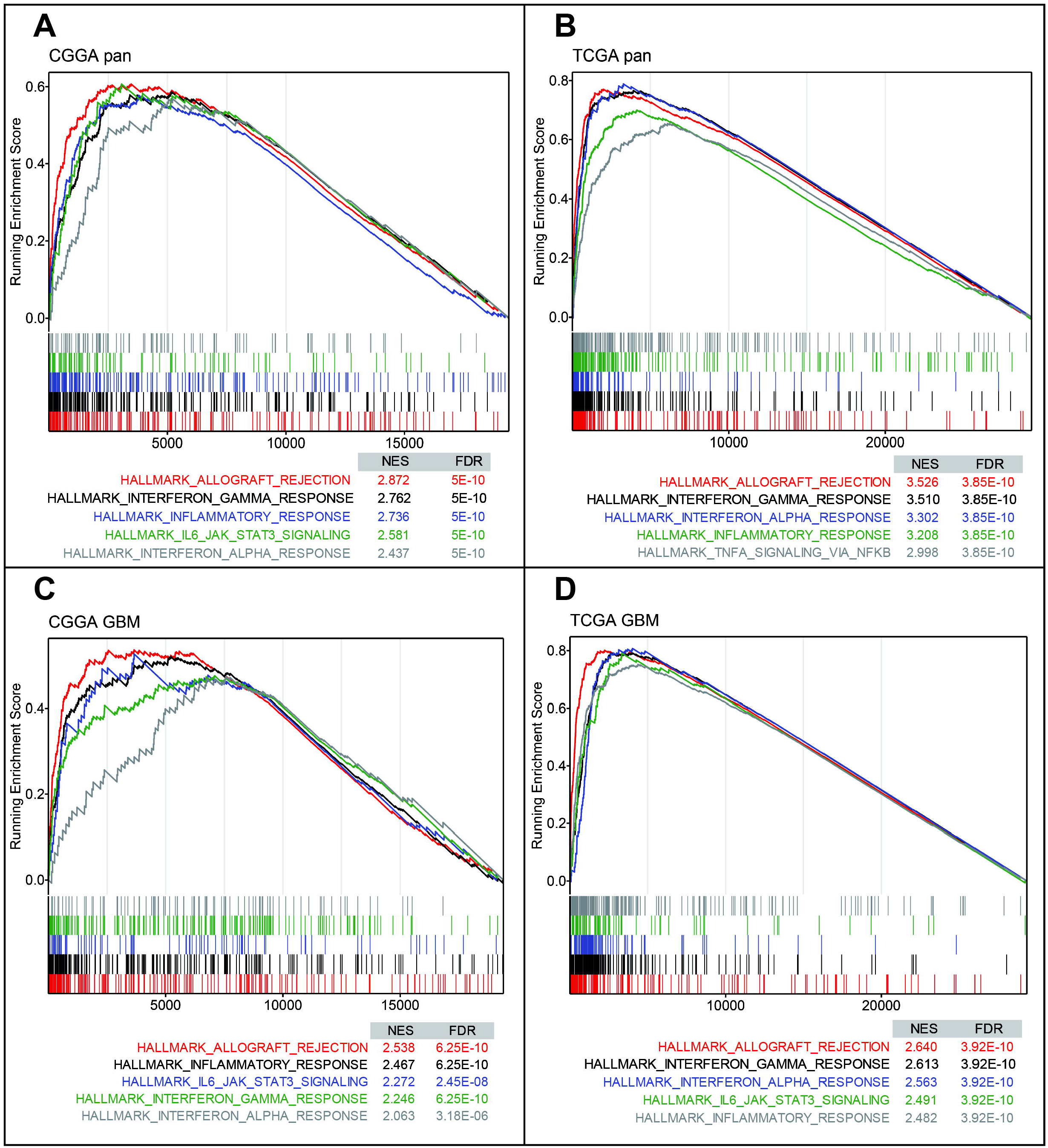
Gene set enrichment analysis (GSEA) for ICOS in pan-glioma (A, B) and glioblastoma (C, D)

### 3.3. ICOS interacts with immune checkpoints in glioma

To further validate the interactions between ICOS and immune response, we performed correlation tests to explore the relationship between ICOS and a series of immune checkpoints, including PD-1, PD-L1, PD-L2, and CTLA-4. Circos plots showed that ICOS level was positively correlated with all of these immune checkpoints both in CGGA dataset and TCGA dataset **(Figures. 4A and 4B)**, exhibiting synergistic interactions of these immune checkpoints in gliomas. Taking GBM as a distinctive group, correlation tests were additionally performed to determine the relationship among these checkpoints in GBM. And it was found that they exhibited more significant correlations with each other (**Figures 4C and 4D**).

**Figure 4.**
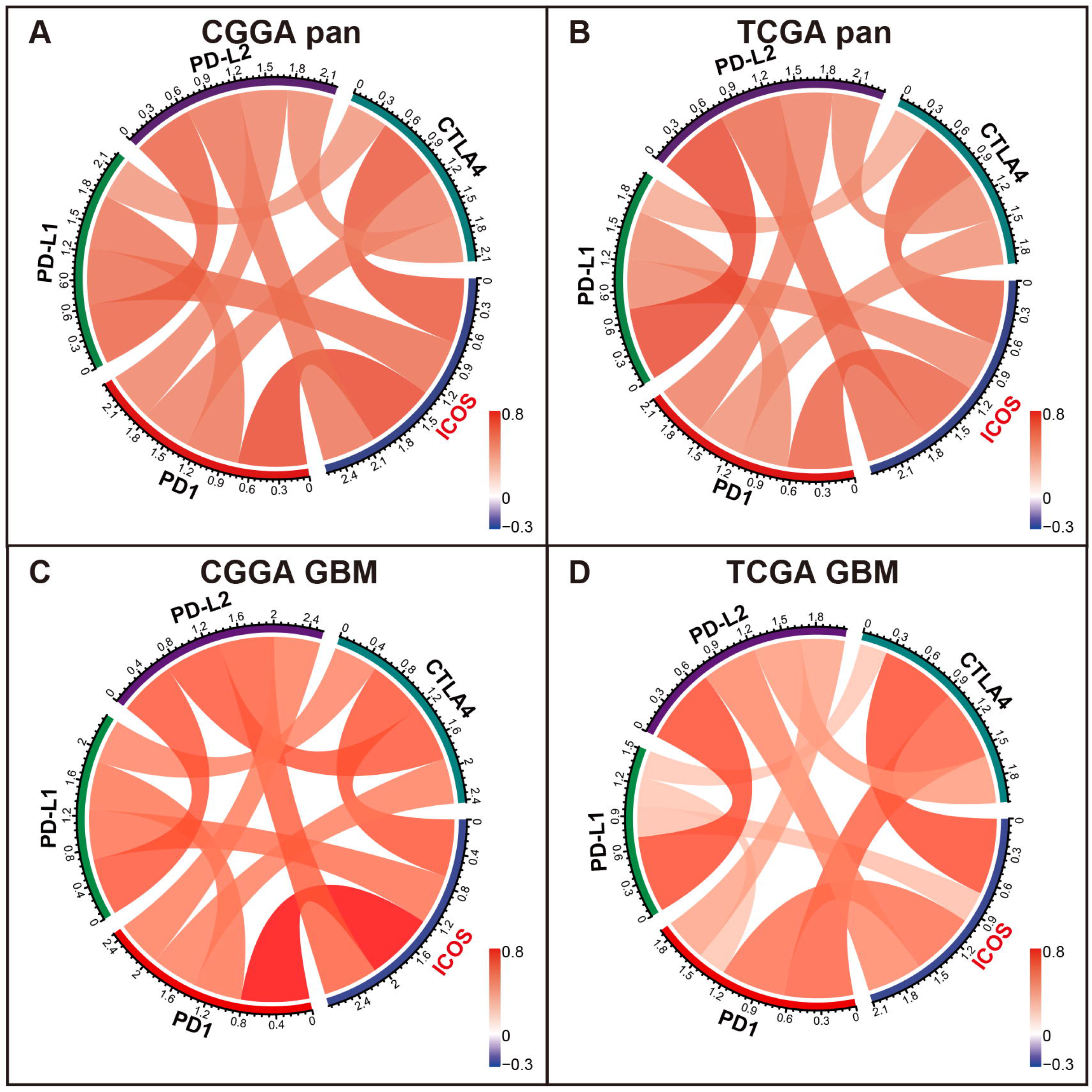
Correlation between ICOS and immune checkpoints in pan-glioma (A, B) and glioblastoma (C, D)

Multiple other checkpoints, such as TIM3, CD28, and IDO1, have been identified as therapeutic targets in preclinical experiments and clinical trials. We additionally examined the correlation between ICOS and these checkpoint members. ICOSLG, the ligand of ICOS, was also included in the analysis. Circos plots demonstrated that ICOS expression showed a robust correlation with **ICOSLG and IDO1 (Figure S2)**. These results reminded us that when glioma acquires resistance to ICOS treatment, we need to be aware of the arsing IDO1.

### 3.4. ICOS-related inflammatory activities

GSVA was performed to identify ICOS-related inflammatory activities. **As shown in Figures 5A and 5B**, ICOS expression revealed a significant correlation with most of the clusters, except for IgG, which specifically represented the immune activities of B cells. Subsequently, correlation tests were performed between the expression levels of seven metagenes and ICOS. As shown in Corrgram plots (**Figures. 5C and 5D**), ICOS was significantly positively correlated with HCK, LCK, MHC-I, MHC-II, STAT1, and Interferon, particularly with LCK, consistent with what we observed in clusters. We observed a similar pattern in GBM of both CGGA and TCGA datasets **(Figure S3)**. These findings enlightened us that ICOS might contribute as a potential immunosuppressor when the function of T cells was activated in gliomas.

**Figure 5.**
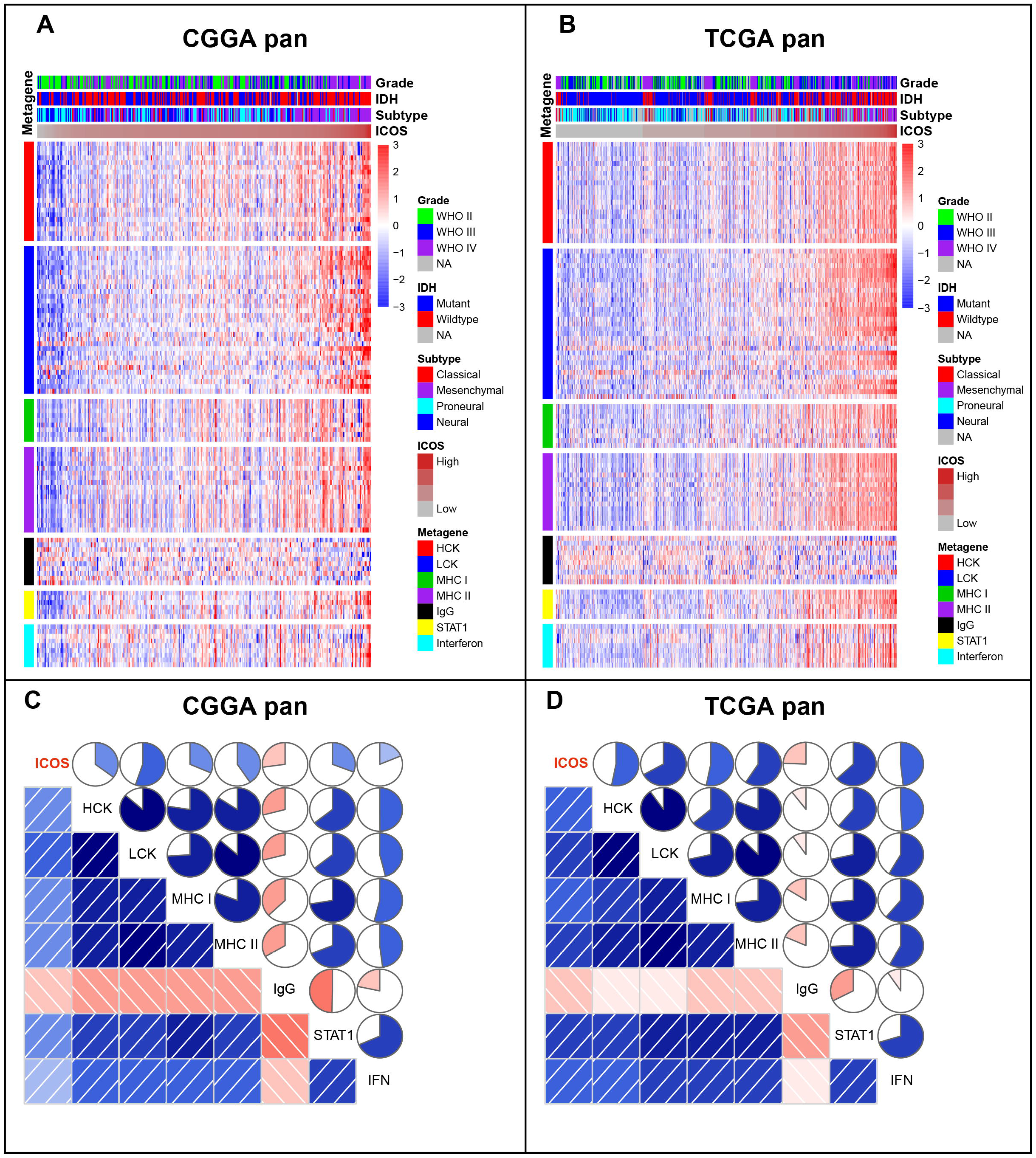
Clusters (A, B) and Gene Sets Variation Analysis (C, D) of ICOS-related inflammatory activities in pan-glioma

### 3.5. Relationship between ICOS and immune cell subpopulations in tumor microenvironment

XCELL analysis(38) was performed to evaluate the immune score, stroma score, microenvironment score, and the cell subpopulations that ICOS might influence in glioma. We found that ICOS was remarkably positively correlated with immune score, stroma score, and microenvironment score of gliomas in both CGGA **(Figure. 6A) and TCGA (Figure. 6B)**. Through cell subpopulation enrichment analysis, ICOS showed a robust correlation with a series of antigen presentation cells, including dendritic cells (DC), monocytes, and macrophages in both cohorts. ICOS was negatively associated with normal brain neurons (**Figure 6**). Noteworthy, the correlation between ICOS and Tregs was relatively not that significant in both cohorts. Overall, these results indicated that gliomas with higher ICOS tend to recruit infiltrating immune cells into the tumor and are significantly associated with alterations in the glioma microenvironment.

**Figure 6.**
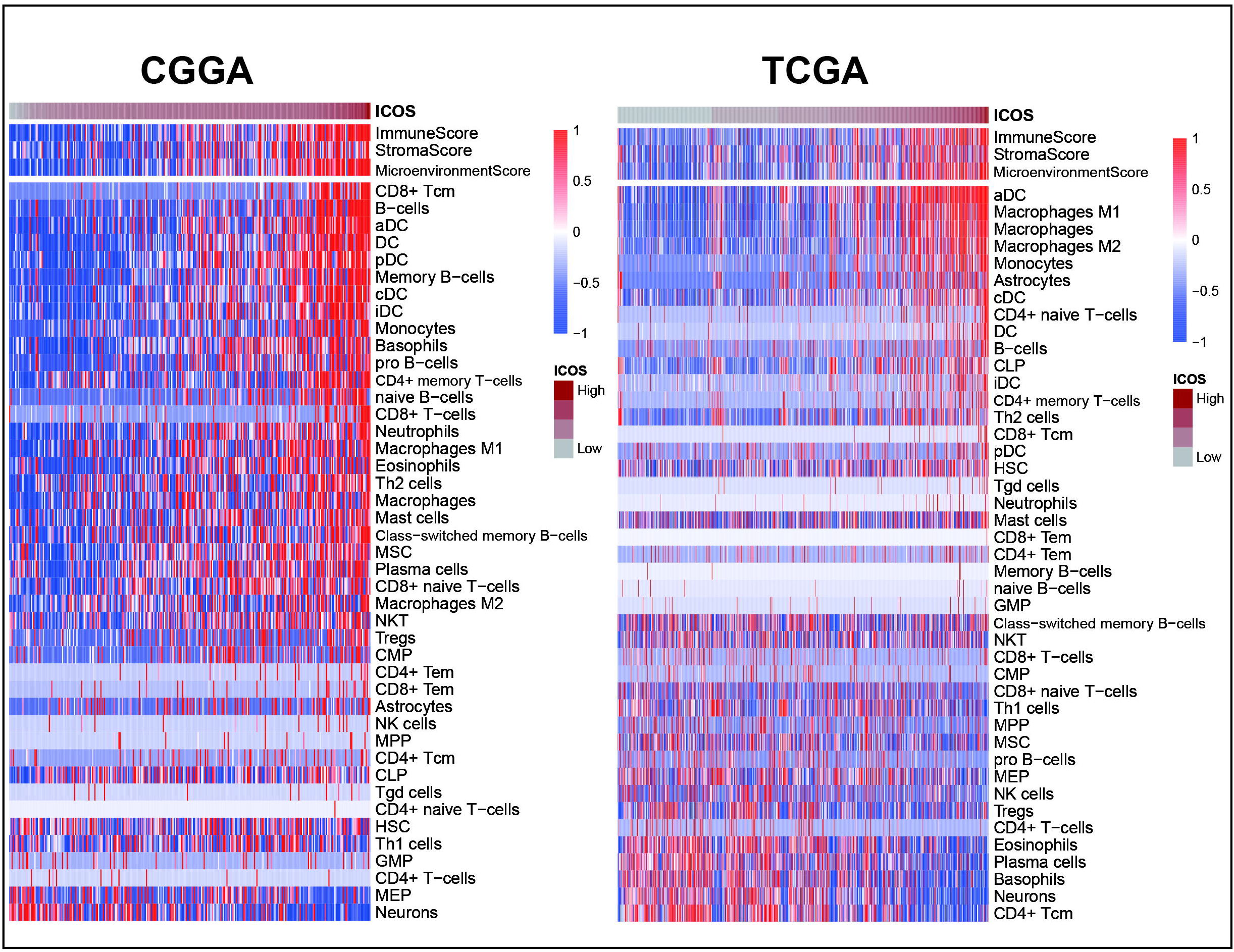
Relationship between ICOS and immune cell subpopulations in pan-glioma

### 3.6. ICOS was mainly expressed by regulatory T cells

To identify the cell types that were highly expressing ICOS, CGGA sc-RNAseq was analyzed. All cells were cut into 5 clusters and were visualized with the UMAP method. Based on the cell markers, cluster 0 overexpressing PDGFRA and cluster 2 overexpressing EGFR could be annotated as glioma cells. Cluster 1 expressing CD68 was annotated as monocyte-macrophage linage, and cluster 4 expressing CD3D was concluded as T cells. Cluster 3, highly expressing MOG, represented a series of normal glial cells. As shown in **Figure 7A**, ICOS was found to be exclusively expressed by T cells (Cluster 4). To explore the T cell subtypes correlated with ICOS upregulation, GSE163108 sc-RNAseq, focusing on T cells of gliomas, was further analyzed. It turned out that ICOS was mainly activated on Tregs (**Figure 7B**), which further validated the immunosuppressive feature of ICOS among gliomas.

**Figure 7.**
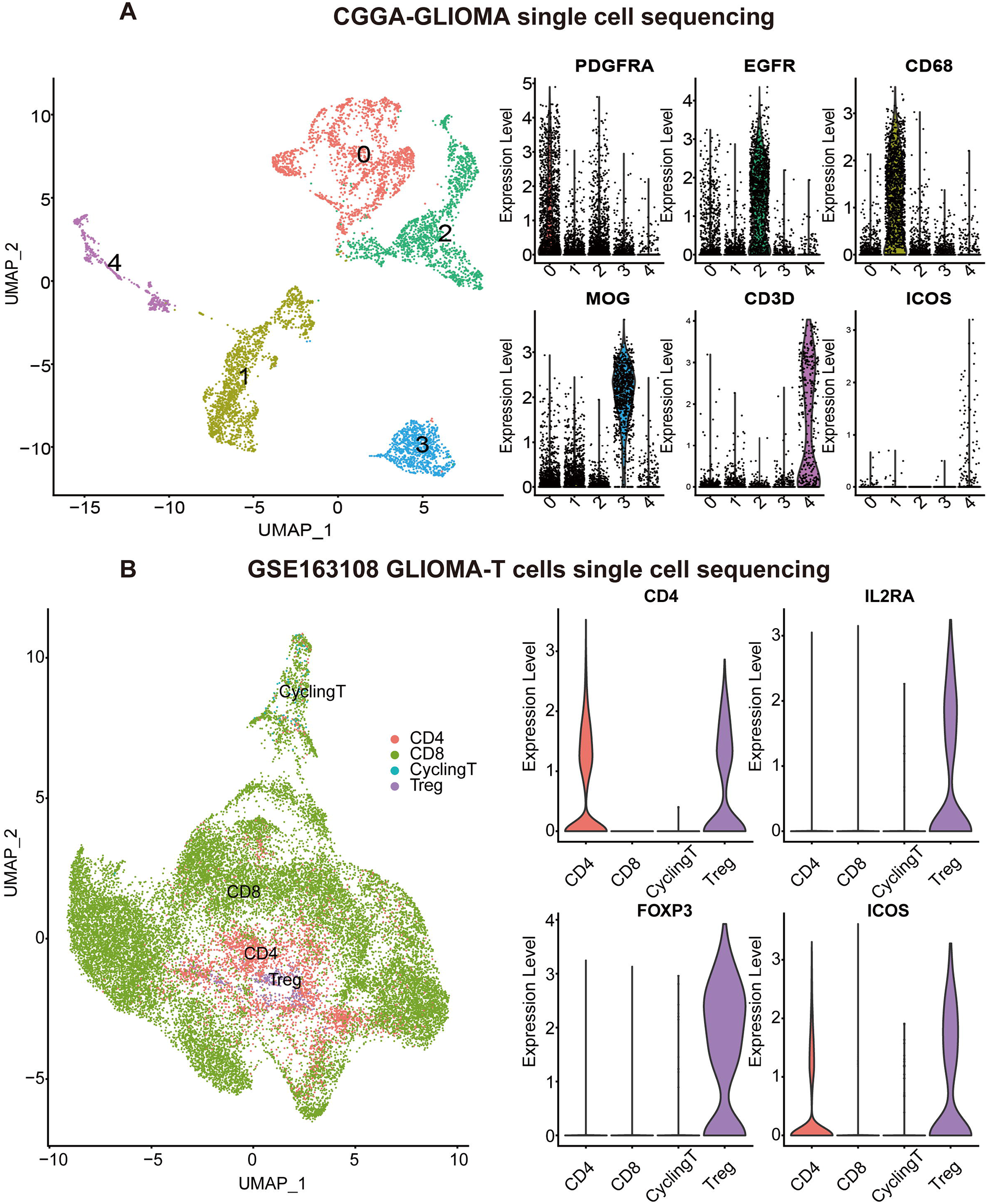
ICOS expression on different cell types based on single-cell RNAseq from CGGA (A) and GSE163108 (B)

### 3.7. ICOS predicts worse survival for gliomas

Kaplan-Meier curves were delineated to explore the prognostic role of ICOS in gliomas. **Figures 8A and 8D** showed that patients with high ICOS levels exhibited significantly worse survival than patients with low ICOS. Similar patterns were observed on the KM curves of LGG **(Figures. 8B and 8E)** and GBM patients **(Figures. 8C and 8F)**. Cox regression analysis was performed to investigate the independent prognostic role of ICOS, together with age, WHO grade, and IDH mutation, and the results revealed that higher ICOS was associated with an unfavorable prognosis independently (CGGA: HR = 1.191, *p* = 0.044; TCGA: HR = 1.151, *p* = 0.005) (**Figures 8G and 8H, Tables S3 and S4**). These findings suggested that ICOS could serve as an independent unfavorable prognostic factor for glioma patients.

**Figure 8.**
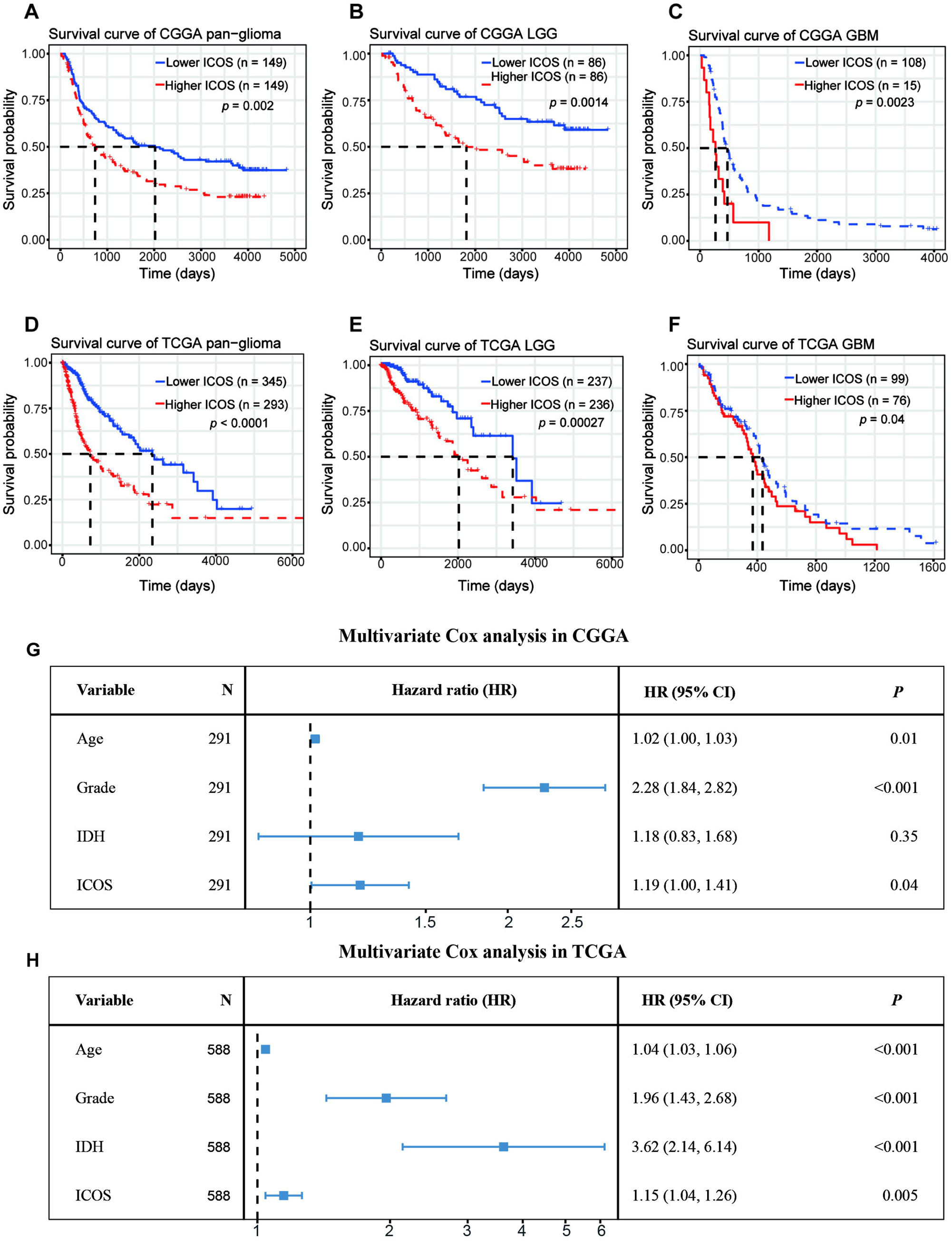
Survival analysis (A-F) and multivariate cox regression analysis (G, H)

## 4. Discussion

Despite numerous efforts to improve prognosis, glioma remains the most common and lethal malignancy in the central nervous system. There is a pressing need to explore novel therapeutic strategies for gliomas. Immunotherapy brings new hope for the treatment of malignancies(40, 41). PD-1 and CTLA4 are two well-known immune checkpoints through which cancers prevent the immune system from recognizing and attacking tumor cells. Anti-PD-1 and anti-CTLA4 can re-activate the anti-tumor response, leading to tumor regression(42, 43). Unfortunately, anti-PD-1 and anti-CTLA4 monotherapy, although an important step in the right direction, was not as effective as it was hoped to be for gliomas(44). Combining existing immune checkpoint blockades and novel targets promises to achieve more success in immunotherapy for glioma patients. As an essential immune costimulatory member, ICOS aroused researchers’ interest because of its dualistic effect during oncogenesis across different tumors.

Our study provides the most comprehensive investigation of the expression profile of ICOS in whole WHO grade gliomas, together with its related molecular signature, clinical significance, and prognostic value. Through analysis of transcriptome expression status of ICOS among 998 glioma patients, we found that higher ICOS expression correlated with higher malignancy of gliomas, based on the fact that ICOS upregulation was remarkably paralleled with much more aggressive characteristics of gliomas, including glioblastoma, IDH-wildtype, and mesenchymal subtype, and could serve as an independent prognosticator for worse survival. These results indicated that ICOS precisely reflected the malignancy of gliomas and could play a pro-oncogenic role in the development and progression of glioma, consistent with the results reported by Iwata et al.(45). Through investigation of expression status and molecule function of the ligand of ICOS (ICOSLG) in glioblastomas, they concluded that ICOSL, via conjugation with ICOS, was associated with more malignancy of glioblastoma. In addition, Our Circosplots also demonstrated the robust interaction between ICOS and ICOSLG in both pan-glioma and GBM, further confirming the pro-oncogenic role of the ICOS/ICOSLG pathway in gliomas.

Although the exact mechanisms of ICOS in the regulation of tumorigenesis and development of glioma are not well understood, some T cell subpopulations might play essential roles in this process. Reportedly, the immunosuppressive effect of ICOS may be accused to the presence of ICOS^+^ regulatory T cells (Treg), accounting for 5-10% of all peripheral CD4^+^ T-cells. Tregs are a subpopulation of T cells that involve in aspects of immunosuppression. Upregulation or activation of Tregs is associated with higher ICOS expression in tumor-infiltrating T cells(29, 45). In this study, through functional analysis of GO and GSEA, we found a consistent result that a higher ICOS level was associated with a more-activated immune response, especially with T cell activity. It was reported that the number of Tregs increased in the glioma microenvironment, and downregulation of Tregs significantly enhanced the effect of immunotherapy(46). ICOS pathway not only promotes Tregs generation via mediating Th17-to-Treg transdifferentiation but also maintains the immunosuppressive effect of Tregs(14). Our sc-RNAseq analysis revealed that ICOS was significantly upregulated in Tregs rather than CD4+ and CD8+ T cells, confirming the immunosuppressive role of ICOS via Tregs activation. While the immune cell subpopulation analysis showed a weak correlation with the relative ratio of Tregs in both cohorts, suggesting that ICOS was more associated with Treg activation, in contrast to a relatively weak effect in promoting Treg generation in gliomas.

Our results demonstrated that ICOS showed a robust correlation with the PD1/PD-L1/PD-L2 pathway, CTLA-4, and indoleamine 2,3-dioxygenase 1 (IDO1), suggesting synergistic interactions among these immune checkpoints. Thus, therapies targeting ICOS/ICOSL pathway in combination with existing immune checkpoint blockades would produce synergistic effects in efficacy, which have been evaluated in preclinical and clinical experiments. To date, four monoclonal-antibodies (mAbs) targeting ICOS are evaluated in clinical trials, among which three are agonistic (GSK3359609, JTX-2011, and KY1044), and one is antagonistic (MEDI-570). The clinical trials mainly focused on anti-ICOS in combination with anti-PD1/PD-L1 or anti-CTLA4 (NCT02904226, NCT02520791, NCT03829501 NCT02723955, and NCT03251924)(47). Zamarin et al.(48)demonstrated that ICOS agonism in tumors enhanced the effectiveness of CTLA-4 blockade in the anti-tumor immune response. Despite activation in Tregs during the treatment, other anti-tumor T cells (CD8^+^ and CD4^+^) were more activated in the tumor microenvironment. Fan et al.(49) also reported a synergistic anti-tumor response of CTLA-4 blockade combined with ICOS stimulation in a mouse cancer model. However, anti-ICOS antagonist mAb also exhibited an anti-tumor effect in some malignancies, including follicular B cell lymphoma(28) and prostate cancer(50), which may be accounted for decreasing in Treg function and proliferation. Thus, considering the remarkable immunosuppressive effect of ICOS, an anti-ICOS antagonist might be more appropriate in glioma; however, this requires a further in-depth understanding of the mechanism of ICOS. Meanwhile, IDO1, a metabolic modulator reported to promote tumors develop immunotherapy resistance (51), has also been identified as an immune target, and the evaluation of anti-IDO1 is undergoing(52). ICOS showed a strong correlation with these molecules, providing more evidence for immunotherapy combination for gliomas.

In addition, the correlation between ICOS expression and seven inflammatory metagenes was further determined. These metagenes represented distinct immune perspectives. The weak correlation between ICOS and IgG was associated with a low abundance of B-linage cells in the CNS. The other 6 ICOS-correlated metagenes mainly reflected the involvement of ICOS in the activities of T-cells (LCK), antigen-presenting cells (HCK, MHC-I, MHC-II), and interferon-response signaling pathway (STAT1, IFN), which was in line with the results above. Moreover, we investigated the association between ICOS and immune score, stroma score, microenvironment score, and immune cell subpopulations. ICOS expression showed a positive correlation with the immune score, stromal score, microenvironment score, and higher-ICOS gliomas tended to recruit multiple infiltrating immune cell types. Given the robust correlation between ICOS and a broad range of immune activities revealed, ICOS is considered to influence the glioma-related immune response profoundly.

The studies performed by Gousias et al.(29), focusing on ICOS+ Tregs, and Iwata et al. (45), characterizing ICOSLG expression in GBM, have done an impressive job and elucidated the vital role of ICOS-pathway in glioma and Tregs. Our study, mainly focusing on transcriptome characterization and clinical significance, extended the research in whole grades of glioma, further highlighted the essential role of ICOS among pan-gliomas, and profoundly expanded the spectrum of research on ICOS. In conclusion, our study reveals the potential of therapies targeting ICOS for glioma immunotherapy. Nevertheless, there is still a long road ahead for further investigating the biological behaviors of ICOS in glioma. Future studies are warranted to evaluate the efficacy and safety of anti-ICOS in combination with other immune checkpoint inhibitors.

## ACKNOWLEDGMENTS

We sincerely thank the Chinese Glioma Genome Atlas (CGGA) team and Cancer Genome Atlas (TCGA) team for their generosity in sharing big data. They have made a substantial dedication to the scientific world and human health. This work was supported by Shenzhen People’s Hospital (SYJCYJ202001).

## AUTHOR CONTRIBUTIONS

Aijun Shan and Fei Shi made contributions to the study conception and design. Jin Wang and Fei Shi participated in data acquisition and data analysis, drafted the manuscript, and revised it critically. All authors have read and approved the final manuscript.

## Supplemental materials

**Figure S1** Gene Ontology analysis of overlapping ICOS-coexpressed genes between CGGA and TCGA datasets. (A) Venn diagram on pan-glioma overlapping genes. (B) GO analysis for pan-glioma overlapping genes. (C) Venn diagram on glioblastoma overlapping genes. (D) GO analysis for glioblastoma overlapping genes.

**Figure S2** Correlation between ICOS and other immune checkpoint members in pan-glioma (A and B) and glioblastoma (C and D).

**Figure S3** Clusters (A, B) and Gene Sets Variation Analysis (C, D) of ICOS-related inflammatory activities in glioblastoma.

**TABLE S1** Patient characteristics in TCGA RNA-seq and CGGA microarray data.

**TABLE S2** Metagenes used for the Gene Sets Variation Analysis.

**TABLE S3** Cox regression analysis of overall survival in CGGA cohort

**TABLE S4** Cox regression analysis of overall survival in TCGA cohort

## REFERENCES

1. Meng X, Zhao Y, Han B, Zha C, Zhang Y, Li Z, et al. Dual functionalized brain-targeting nanoinhibitors restrain temozolomide-resistant glioma via attenuating EGFR and MET signaling pathways. Nat Commun. 2020;11(1):594.

2. Wang X, Lu S, He C, Wang C, Wang L, Piao M, et al. RSL3 induced autophagic death in glioma cells via causing glycolysis dysfunction. Biochem Biophys Res Commun. 2019;518(3):590–7.

3. Wei J, Ouyang X, Tang Y, Li H, Wang B, Ye Y, et al. ER-stressed MSC displayed more effective immunomodulation in RA CD4(+)CXCR5(+)ICOS(+) follicular helper-like T cells through higher PGE2 binding with EP2/EP4. Mod Rheumatol. 2019:1–8.

4. Hutloff A, Dittrich AM, Beier KC, Eljaschewitsch B, Kraft R, Anagnostopoulos I, et al. ICOS is an inducible T-cell co-stimulator structurally and functionally related to CD28. Nature. 1999;397(6716):263–6.

5. Wikenheiser DJ, Stumhofer JS. ICOS Co-Stimulation: Friend or Foe? Front Immunol. 2016;7:304.

6. Schepp J, Chou J, Skrabl-Baumgartner A, Arkwright PD, Engelhardt KR, Hambleton S, et al. 14 Years after Discovery: Clinical Follow-up on 15 Patients with Inducible Co-Stimulator Deficiency. Front Immunol. 2017;8:964.

7. Pratama A, Srivastava M, Williams NJ, Papa I, Lee SK, Dinh XT, et al. MicroRNA-146a regulates ICOS-ICOSL signalling to limit accumulation of T follicular helper cells and germinal centres. Nat Commun. 2015;6:6436.

8. Stone EL, Pepper M, Katayama CD, Kerdiles YM, Lai CY, Emslie E, et al. ICOS coreceptor signaling inactivates the transcription factor FOXO1 to promote Tfh cell differentiation. Immunity. 2015;42(2):239–51.

9. Mesturini R, Nicola S, Chiocchetti A, Bernardone IS, Castelli L, Bensi T, et al. ICOS cooperates with CD28, IL-2, and IFN-gamma and modulates activation of human naive CD4+ T cells. Eur J Immunol. 2006;36(10):2601–12.

10. Mesturini R, Gigliotti CL, Orilieri E, Cappellano G, Soluri MF, Boggio E, et al. Differential induction of IL-17, IL-10, and IL-9 in human T helper cells by B7h and B7.1. Cytokine. 2013;64(1):322–30.

11. Wallin JJ, Liang L, Bakardjiev A, Sha WC. Enhancement of CD8+ T cell responses by ICOS/B7h costimulation. J Immunol. 2001;167(1):132–9.

12. Nelson MH, Kundimi S, Bowers JS, Rogers CE, Huff LW, Schwartz KM, et al. The inducible costimulator augments Tc17 cell responses to self and tumor tissue. J Immunol. 2015;194(4):1737–47.

13. Herman S, Zurgil N, Langevitz P, Ehrenfeld M, Deutsch M. The immunosuppressive effect of methotrexate in active rheumatoid arthritis patients vs. its stimulatory effect in nonactive patients, as indicated by cytometric measurements of CD4+ T cell subpopulations. Immunol Invest. 2004;33(3):351–62.

14. Downs-Canner S, Berkey S, Delgoffe GM, Edwards RP, Curiel T, Odunsi K, et al. Suppressive IL-17A(+)Foxp3(+) and ex-Th17 IL-17A(neg)Foxp3(+) Treg cells are a source of tumour-associated Treg cells. Nat Commun. 2017;8:14649.

15. Edwards J, Tasker A, Pires da Silva I, Quek C, Batten M, Ferguson A, et al. Prevalence and Cellular Distribution of Novel Immune Checkpoint Targets Across Longitudinal Specimens in Treatment-naive Melanoma Patients: Implications for Clinical Trials. Clin Cancer Res. 2019;25(11):3247–58.

16. Strauss L, Bergmann C, Szczepanski MJ, Lang S, Kirkwood JM, Whiteside TL. Expression of ICOS on human melanoma-infiltrating CD4+CD25highFoxp3+ T regulatory cells: implications and impact on tumor-mediated immune suppression. J Immunol. 2008;180(5):2967–80.

17. Bogunovic D, O’Neill DW, Belitskaya-Levy I, Vacic V, Yu YL, Adams S, et al. Immune profile and mitotic index of metastatic melanoma lesions enhance clinical staging in predicting patient survival. Proc Natl Acad Sci U S A. 2009;106(48):20429–34.

18. Nagase H, Takeoka T, Urakawa S, Morimoto-Okazawa A, Kawashima A, Iwahori K, et al. ICOS(+) Foxp3(+) TILs in gastric cancer are prognostic markers and effector regulatory T cells associated with Helicobacter pylori. Int J Cancer. 2017;140(3):686–95.

19. Huang XM, Liu XS, Lin XK, Yu H, Sun JY, Liu XK, et al. Role of plasmacytoid dendritic cells and inducible costimulator-positive regulatory T cells in the immunosuppression microenvironment of gastric cancer. Cancer Sci. 2014;105(2):150–8.

20. Pedroza-Gonzalez A, Zhou G, Vargas-Mendez E, Boor PP, Mancham S, Verhoef C, et al. Tumor-infiltrating plasmacytoid dendritic cells promote immunosuppression by Tr1 cells in human liver tumors. Oncoimmunology. 2015;4(6):e1008355.

21. Toker A, Nguyen LT, Stone SC, Yang SYC, Katz SR, Shaw PA, et al. Regulatory T Cells in Ovarian Cancer Are Characterized by a Highly Activated Phenotype Distinct from that in Melanoma. Clin Cancer Res. 2018;24(22):5685–96.

22. Conrad C, Gregorio J, Wang YH, Ito T, Meller S, Hanabuchi S, et al. Plasmacytoid dendritic cells promote immunosuppression in ovarian cancer via ICOS costimulation of Foxp3(+) T-regulatory cells. Cancer Res. 2012;72(20):5240–9.

23. Faget J, Bendriss-Vermare N, Gobert M, Durand I, Olive D, Biota C, et al. ICOS-ligand expression on plasmacytoid dendritic cells supports breast cancer progression by promoting the accumulation of immunosuppressive CD4+ T cells. Cancer Res. 2012;72(23):6130–41.

24. Faget J, Sisirak V, Blay JY, Caux C, Bendriss-Vermare N, Menetrier-Caux C. ICOS is associated with poor prognosis in breast cancer as it promotes the amplification of immunosuppressive CD4(+) T cells by plasmacytoid dendritic cells. Oncoimmunology. 2013;2(3):e23185.

25. Giraldo NA, Becht E, Vano Y, Petitprez F, Lacroix L, Validire P, et al. Tumor-Infiltrating and Peripheral Blood T-cell Immunophenotypes Predict Early Relapse in Localized Clear Cell Renal Cell Carcinoma. Clin Cancer Res. 2017;23(15):4416–28.

26. Zhang Y, Luo Y, Qin SL, Mu YF, Qi Y, Yu MH, et al. The clinical impact of ICOS signal in colorectal cancer patients. Oncoimmunology. 2016;5(5):e1141857.

27. Lee H, Kim JH, Yang SY, Kong J, Oh M, Jeong DH, et al. Peripheral blood gene expression of B7 and CD28 family members associated with tumor progression and microscopic lymphovascular invasion in colon cancer patients. J Cancer Res Clin Oncol. 2010;136(9):1445–52.

28. Le KS, Thibult ML, Just-Landi S, Pastor S, Gondois-Rey F, Granjeaud S, et al. Follicular B Lymphomas Generate Regulatory T Cells via the ICOS/ICOSL Pathway and Are Susceptible to Treatment by Anti-ICOS/ICOSL Therapy. Cancer Res. 2016;76(16):4648–60.

29. Gousias K, von Ruecker A, Voulgari P, Simon M. Phenotypical analysis, relation to malignancy and prognostic relevance of ICOS+T regulatory and dendritic cells in patients with gliomas. J Neuroimmunol. 2013;264(1-2):84–90.

30. Zhao Z, Zhang KN, Wang Q, Li G, Zeng F, Zhang Y, et al. Chinese Glioma Genome Atlas (CGGA): A Comprehensive Resource with Functional Genomic Data from Chinese Glioma Patients. Genomics Proteomics Bioinformatics. 2021;19(1):1–12.

31. Cancer Genome Atlas Research N, Weinstein JN, Collisson EA, Mills GB, Shaw KR, Ozenberger BA, et al. The Cancer Genome Atlas Pan-Cancer analysis project. Nat Genet. 2013;45(10):1113–20.

32. Yu K, Hu Y, Wu F, Guo Q, Qian Z, Hu W, et al. Surveying brain tumor heterogeneity by single-cell RNA-sequencing of multi-sector biopsies. Natl Sci Rev. 2020;7(8):1306–18.

33. Mathewson ND, Ashenberg O, Tirosh I, Gritsch S, Perez EM, Marx S, et al. Inhibitory CD161 receptor identified in glioma-infiltrating T cells by single-cell analysis. Cell. 2021;184(5):1281–98 e26.

34. Sherman BT, Hao M, Qiu J, Jiao X, Baseler MW, Lane HC, et al. DAVID: a web server for functional enrichment analysis and functional annotation of gene lists (2021 update). Nucleic Acids Res. 2022.

35. Subramanian A, Tamayo P, Mootha VK, Mukherjee S, Ebert BL, Gillette MA, et al. Gene set enrichment analysis: a knowledge-based approach for interpreting genome-wide expression profiles. Proc Natl Acad Sci U S A. 2005;102(43):15545–50.

36. Rody A, Holtrich U, Pusztai L, Liedtke C, Gaetje R, Ruckhaeberle E, et al. T-cell metagene predicts a favorable prognosis in estrogen receptor-negative and HER2-positive breast cancers. Breast Cancer Res. 2009;11(2):R15.

37. Hanzelmann S, Castelo R, Guinney J. GSVA: gene set variation analysis for microarray and RNA-seq data. BMC Bioinformatics. 2013;14:7.

38. Aran D, Hu Z, Butte AJ. xCell: digitally portraying the tissue cellular heterogeneity landscape. Genome Biol. 2017;18(1):220.

39. Yu G, Wang LG, Han Y, He QY. clusterProfiler: an R package for comparing biological themes among gene clusters. OMICS. 2012;16(5):284–7.

40. Li G, Wang Z, Zhang C, Liu X, Cai J, Wang Z, et al. Molecular and clinical characterization of TIM-3 in glioma through 1,024 samples. Oncoimmunology. 2017;6(8):e1328339.

41. Wang ZL, Li GZ, Wang QW, Bao ZS, Wang Z, Zhang CB, et al. PD-L2 expression is correlated with the molecular and clinical features of glioma, and acts as an unfavorable prognostic factor. Oncoimmunology. 2019;8(2):e1541535.

42. Liu S, Wang Z, Wang Y, Fan X, Zhang C, Ma W, et al. PD-1 related transcriptome profile and clinical outcome in diffuse gliomas. Oncoimmunology. 2018;7(2):e1382792.

43. Liu F, Huang J, Liu X, Cheng Q, Luo C, Liu Z. CTLA-4 correlates with immune and clinical characteristics of glioma. Cancer Cell Int. 2020;20:7.

44. Brown CE, Alizadeh D, Starr R, Weng L, Wagner JR, Naranjo A, et al. Regression of Glioblastoma after Chimeric Antigen Receptor T-Cell Therapy. N Engl J Med. 2016;375(26):2561–9.

45. Iwata R, Hyoung Lee J, Hayashi M, Dianzani U, Ofune K, Maruyama M, et al. ICOSLG-mediated regulatory T-cell expansion and IL-10 production promote progression of glioblastoma. Neuro Oncol. 2020;22(3):333–44.

46. Vandenberk L, Van Gool SW. Treg infiltration in glioma: a hurdle for antiglioma immunotherapy. Immunotherapy. 2012;4(7):675–8.

47. Morad G, Helmink BA, Sharma P, Wargo JA. Hallmarks of response, resistance, and toxicity to immune checkpoint blockade. Cell. 2021;184(21):5309–37.

48. Zamarin D, Holmgaard RB, Ricca J, Plitt T, Palese P, Sharma P, et al. Intratumoral modulation of the inducible co-stimulator ICOS by recombinant oncolytic virus promotes systemic anti-tumour immunity. Nat Commun. 2017;8:14340.

49. Fan X, Quezada SA, Sepulveda MA, Sharma P, Allison JP. Engagement of the ICOS pathway markedly enhances efficacy of CTLA-4 blockade in cancer immunotherapy. J Exp Med. 2014;211(4):715–25.

50. Mo L, Chen Q, Zhang X, Shi X, Wei L, Zheng D, et al. Depletion of regulatory T cells by anti-ICOS antibody enhances anti-tumor immunity of tumor cell vaccine in prostate cancer. Vaccine. 2017;35(43):5932–8.

51. Munn DH, Mellor AL. Indoleamine 2,3-dioxygenase and tumor-induced tolerance. J Clin Invest. 2007;117(5):1147–54.

52. Guo Y, Liu Y, Wu W, Ling D, Zhang Q, Zhao P, et al. Indoleamine 2,3-dioxygenase (Ido) inhibitors and their nanomedicines for cancer immunotherapy. Biomaterials. 2021;276:121018.

