## Supplemental Table 1 for "Transcriptome profile and clinical characterization of ICOS expression in brain gliomas"

**Supplemental Table 1. Patient characteristics in TCGA RNA-seq and CGGA microarray data.**

| **Characteristics** | **TCGA RNA-seq (n=697)** | **CGGA microarray (n=301)** |
| --- | --- | --- |
| **Gender** |  |  |
| male | 370 | 180 |
| female | 271 | 121 |
| NA | 56 | 0 |
| **Age (year)** | 47 ± 15 | 42 ± 12 |
| **Tumor subtype** |  |  |
| Classical | 90 | 23 |
| Mesenchymal | 104 | 111 |
| Proneural | 248 | 86 |
| Neural | 115 | 81 |
| NA | 140 | 0 |
| **WHO grade** |  |  |
| Grade II | 226 | 122 |
| Grade III | 249 | 51 |
| Grade IV | 167 | 128 |
| NA | 55 | 0 |
| **Karnofsky Performance Score** | 84 ± 14 | NA |
| **IDH mutation status** |  |  |
| Mut | 442 | 134 |
| WT | 245 | 165 |
| NA | 10 | 2 |
| **1p/19q Codeletion status** |  |  |
| Codeletion | 181 | 16 |
| Non-codeletion | 491 | 76 |
| NA | 25 | 209 |
| **MGMT promoter status** |  |  |
| Methylated | 461 | 99 |
| Unmethylated | 162 | 187 |
| NA | 74 | 15 |

NA: Not Available; KPS: Karnofsky Performance Score; MGMT: O^6^-Methylguanine Methyltransferase
