## Supplemental Table 2 for "Transcriptome profile and clinical characterization of ICOS expression in brain gliomas"

**Supplemental Table 2 Metagenes used for the Gene Sets Variation Analysis**

| **Metagenes** | **Genes** |
| --- | --- |
| HCK | C1QB |
| HCK | C1QA |
| HCK | AIF1 |
| HCK | LST1 |
| HCK | DOCK2 |
| HCK | LAPTM5 |
| HCK | TYROBP |
| HCK | MS4A4A |
| HCK | MS4A6A |
| HCK | CD163 |
| HCK | ITGB2 |
| HCK | SLC7A7 |
| HCK | LAIR1 |
| HCK | HCK |
| HCK | TFEC |
| HCK | IFI30 |
| HCK | MNDA |
| HCK | FCER1G |
| HCK | RNASE6 |
| HCK | SLCO2B1 |
| HCK | CCR1 |
| IgG | IGSF8 |
| IgG | ISLR2 |
| IgG | IGSF21 |
| IgG | IGSF1 |
| IgG | IGSF22 |
| IgG | IGDCC3 |
| IgG | IGHD |
| IgG | IGSF11 |
| IgG | IGSF5 |
| IgG | IGSF6 |
| Interferon | IFIT1 |
| Interferon | IFIT3 |
| Interferon | IFI44L |
| Interferon | OAS3 |
| Interferon | MX1 |
| Interferon | RSAD2 |
| Interferon | IFI44 |
| Interferon | OAS2 |
| Interferon | OAS1 |
| LCK | CD2 |
| LCK | GZMK |
| LCK | GZMA |
| LCK | CD3D |
| LCK | CD53 |
| LCK | LCK |
| LCK | ARHGAP15 |
| LCK | CCL5 |
| LCK | GMFG |
| LCK | SELL |
| LCK | STAT4 |
| LCK | SAMSN1 |
| LCK | RAC2 |
| LCK | HCLS1 |
| LCK | CCR7 |
| LCK | PIK3CD |
| LCK | CORO1A |
| LCK | CD48 |
| LCK | IL2RG |
| LCK | SH2D1A |
| LCK | SLAMF1 |
| LCK | IL7R |
| LCK | INPP5D |
| LCK | KLRK1 |
| LCK | FGL2 |
| LCK | IRF8 |
| LCK | SELPLG |
| LCK | IL10RA |
| LCK | SLA |
| LCK | CCR2 |
| LCK | CSF2RB |
| MHC_I | HLA-E |
| MHC_I | HLA-H |
| MHC_I | HLA-B |
| MHC_I | HLA-J |
| MHC_I | HLA-F |
| MHC_I | HLA-G |
| MHC_I | HLA-A |
| MHC_I | HLA-C |
| MHC_I | HLA-L |
| MHC_II | HLA-DRB1 |
| MHC_II | HLA-DRB5 |
| MHC_II | HLA-DRB3 |
| MHC_II | HLA-DPA1 |
| MHC_II | HLA-DRA |
| MHC_II | HLA-DQA1 |
| MHC_II | HLA-DQA2 |
| MHC_II | HLA-DMA |
| MHC_II | HLA-DOA |
| MHC_II | HLA-DRB4 |
| MHC_II | HLA-DMB |
| MHC_II | HLA-DQB1 |
| MHC_II | HLA-DPB1 |
| MHC_II | HLA-DQB2 |
| MHC_II | CD74 |
| MHC_II | PTPRC |
| MHC_II | HLA-DOB |
| MHC_II | HLA-DPB2 |
| STAT1 | TAP1 |
| STAT1 | STAT1 |
| STAT1 | CXCL10 |
| STAT1 | CXCL11 |
| STAT1 | GBP1 |
| STAT1 | CXCL9 |
