## Supplemental Table 3 for "Transcriptome profile and clinical characterization of ICOS expression in brain gliomas"

**TABLE S3.** Cox regression analysis of overall survival in CGGA cohort

|  |  | **Univariate** | | |  | **Multivariate** | | |
| --- | --- | --- | --- | --- | --- | --- | --- | --- |
| **Covariates** |  | **HR** | **95% CI** | ***P* value** |  | **HR** | **95% CI** | ***P* value** |
| Age (years) |  | 1.041 | 1.027-1.055 | **0.000** |  | 1.017 | 1.004-1.031 | **0.012** |
| WHO Grade (II-IV) |  | 2.670 | 2.221-3.210 | **0.000** |  | 2.275 | 1.838-2.816 | **0.000** |
| IDH (ref. Mutant type) |  | 2.541 | 1.869-3.454 | **0.000** |  | 1.184 | 0.834-1.682 | 0.345 |
| ICOS expression |  | 1.598 | 1.367-1.869 | **0.000** |  | 1.191 | 1.004-1.413 | **0.044** |
