## Supplemental Table 4 for "Transcriptome profile and clinical characterization of ICOS expression in brain gliomas"

**TABLE S4.** Cox regression analysis of overall survival in TCGA cohort

|  |  | **Univariate** | | |  | **Multivariate** | | |
| --- | --- | --- | --- | --- | --- | --- | --- | --- |
| **Covariates** |  | **HR** | **95%CI** | ***P* value** |  | **HR** | **95%CI** | ***P* value** |
| Age (year) |  | 1.075 | 1.062-1.087 | **0.000** |  | 1.043 | 1.028-1.059 | **0.000** |
| WHO Grade (II-IV) |  | 5.057 | 3.915-6.532 | **0.000** |  | 1.963 | 1.431-2.680 | **0.000** |
| IDH status (ref. Mutant type) |  | 13.101 | 8.962-19.150 | **0.000** |  | 3.622 | 2.141-6.138 | **0.000** |
| ICOS expression |  | 1.545 | 1.437-1.611 | **0.000** |  | 1.151 | 1.040-1.261 | **0.005** |
