## Supplementary figures and images for "Transcriptome profile and clinical characterization of ICOS expression in brain gliomas"

### Supplemental Figure 1

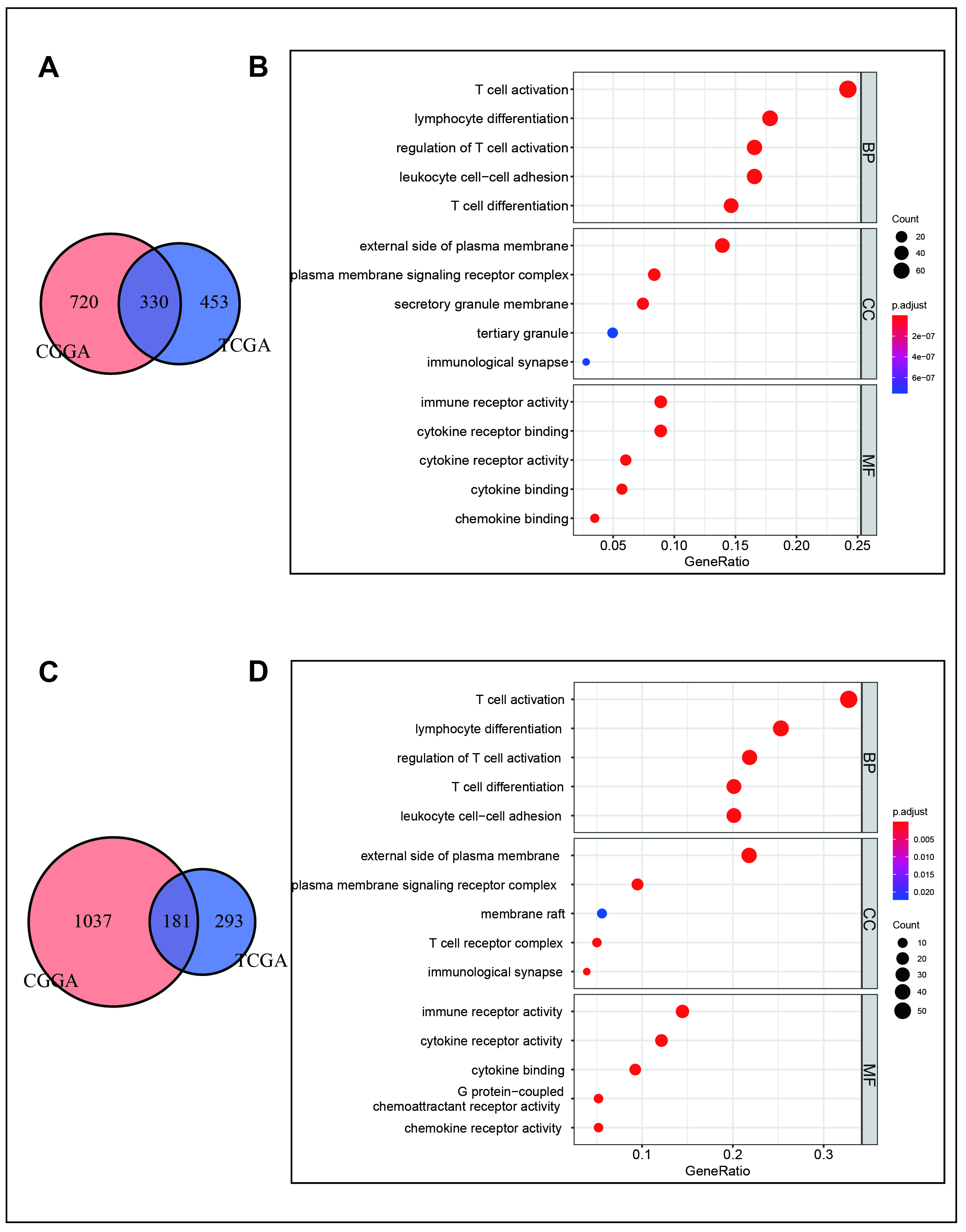

### Supplemental Figure 2

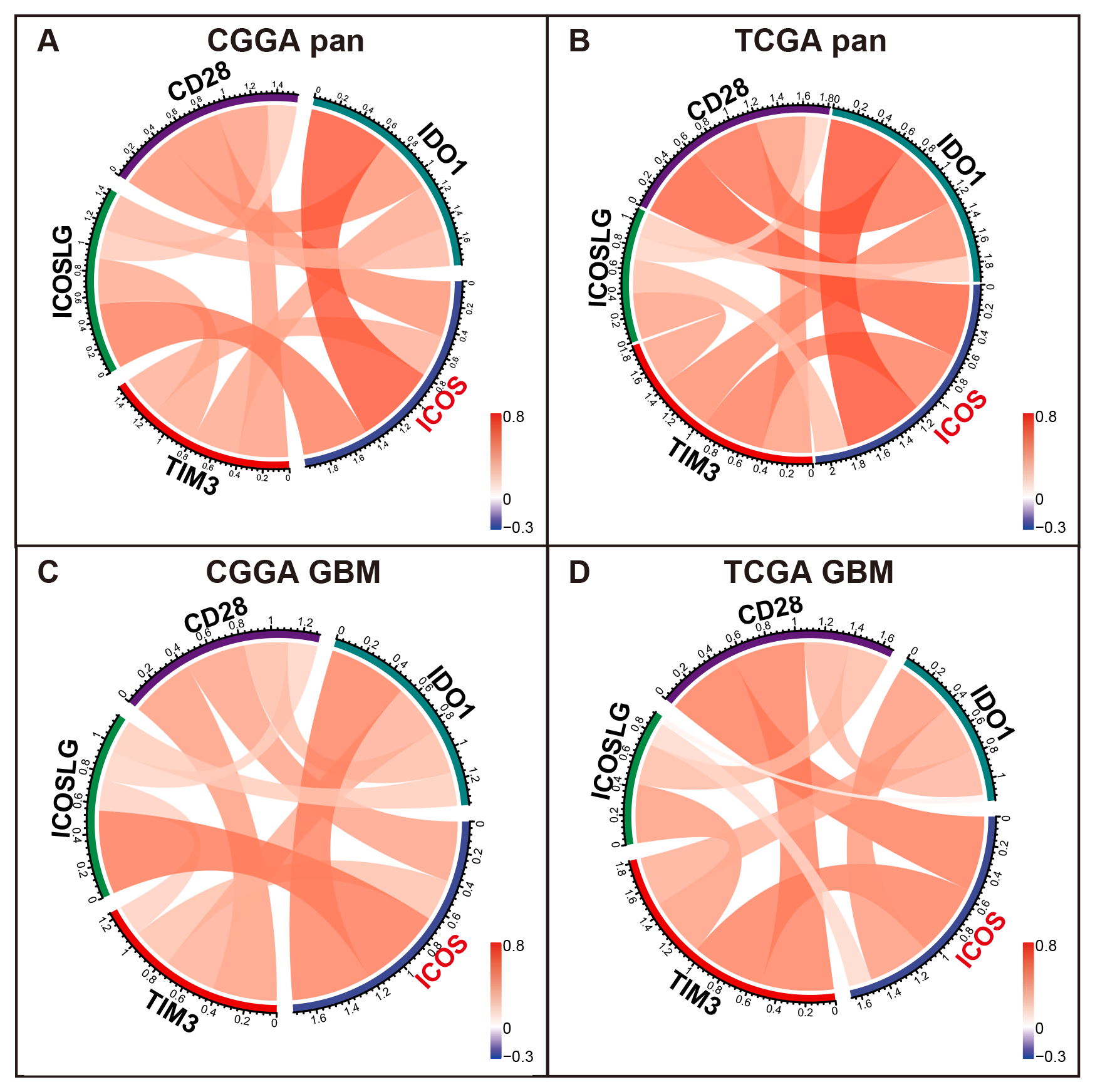

### Supplemental Figure 3

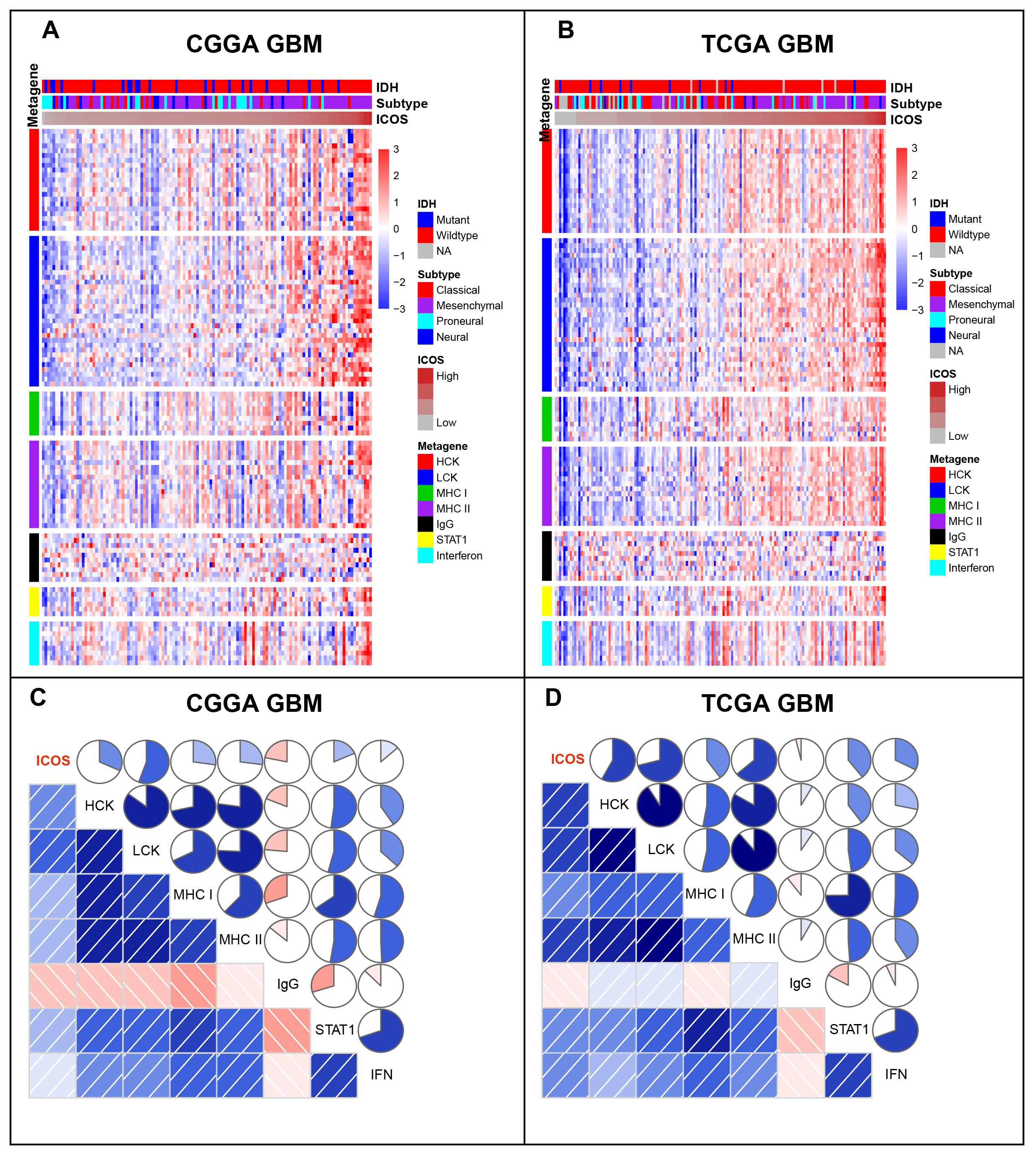
